# Holotomography: refractive index as an intrinsic imaging contrast for 3-D label-free live cell imaging

**DOI:** 10.1101/106328

**Authors:** Doyeon Kim, SangYun Lee, Moosung Lee, JunTaek Oh, Su-A Yang, YongKeun Park

## Abstract

Live cell imaging provides essential information in the investigation of cell biology and related pathophysiology. Refractive index (RI) can serve as intrinsic optical imaging contrast for 3-D label-free and quantitative live cell imaging, and provide invaluable information to understand various dynamics of cells and tissues for the study of numerous fields. Recently significant advances have been made in imaging methods and analysis approaches utilizing RI, which are now being transferred to biological and medical research fields, providing novel approaches to investigate the pathophysiology of cells. To provide insight how RI can be used as an imaging contrast for imaging of biological specimens, here we provide the basic principle of RI-based imaging techniques and summarize recent progress on applications, ranging from microbiology, hematology, infectious diseases, hematology, and histopathology.

## Introduction

Optical images of biological cells and tissues provide invaluable information on the pathophysiology of diseases. Visual diagnosis of blood cells is essential for the diagnosis of various infectious diseases associated with red blood cells (Kaushansky et al. 2016). In pathology and cytology, tissue biopsy and morphological examination of cells, such as checking for abnormally shaped nuclei in the Papanicolaou test, is an essential step for cancer diagnosis (Koss 1989).

The medical diagnostic capabilities of various diseases have evolved with advances in optical imaging technology. In the 17th century, Robert Hooke first observed cork cells using his microscope. Since then, various types of microscopes have been developed and likewise the ability to investigate disease-related cellular, and cellular structures have dramatically improved in recent decades. For example, the invention of phase contrast and differential interference contrast microscopes in the mid 20^th^ century have accelerated the studies in the field of microbiology and cell biology, because these interferometric microscopy techniques allowed effective visualization of transparent biological cells (Allen et al. 1969; Zernike 1942). The development of fluorescent proteins and fluorescence microscopy have also enabled specific labeling of target molecules or proteins. This breakthrough in technology thus has opened a new era for molecular biology.

Moreover, various super-resolution microscope techniques have broken the barrier of diffraction-limited optical resolution. The resolution limit of the optical microscope has been extended to the nanometer scale, enabling investigation of biological phenomena at the single molecule scale (Hell 2007; Huang et al. 2009). More recently, researchers have used adaptive optical approaches for *in vivo* imaging of biological cells or tissues (Ji et al. 2010; Park et al. 2017b; Yu et al. 2014; Yu et al. 2015).

Over the past several decades, fluorescent protein technology has been widely used to locate specific target molecules and proteins in cells using the molecular specificity of the probe (Specht et al. 2017). This allows effective visualization of specific targets in cells and tissues with very high imaging contrast (Lichtman and Conchello 2005). When combined with the fluorescence correlation spectroscopy technique, the fluorescent probe can also provide information about the physical and chemical information of the surrounding medium (Thompson 2002). In addition, when combined with the Foster resonance energy transfer technique, the intermolecular distance can be accurately measured in the nanometer scale (Roy et al. 2008).

However, the use of fluorescent probes in biological imaging inevitably causes several limitations. It is important to note that fluorescence technology uses exogenous fluorescent molecules as a secondary imaging contrast. Expression or binding of a target molecule of a fluorescent probe in a cell generates various problems (Fei et al. 2011; Hoebe et al. 2007). First, the introduction of an exogenous marker into a cell can affect the intrinsic physiology of the cell, because of possible photodamage and phototoxicity caused by the fluorescent molecule. This issue becomes even more severe when experimenting with neurons or stem cells because these cells are more sensitive to changes in the environment (Braydich-Stolle et al. 2005; Millet et al. 2007). Second, long-term cell imaging may be limited when using fluorescent technologies due to photobleaching of probes (Hoebe et al. 2007). Common fluorescent probes cannot produce strong, continuous fluorescence signals. Most probes photobleach after some time, which means the probes irreversibly lose its fluorescence.

Thus maximum period for long-term imaging of live cells is limited to the photobleaching period of fluorescent probes. Third, most fluorescence technologies do not provide quantitative information. The use of fluorescent probes only provides information of the location of the target molecule, but it does not provide quantified information of the mass or concentration of the target molecule.

To complement the limitation caused by the use of exogenous imaging contrast, the use of refractive index (RI), as an intrinsic optical parameter has been exploited recently. All materials have unique RI value, which is correlated with the electrical permittivity of the material. RI is the ratio of the speed of light passing through the specific material to that passing in the vacuum. Conventional phase contrast or differential interference microscopy uses RI values as optical imaging contrast. However, their imaging systems do not provide a one-to-one quantitative mapping of the information about RI distributions in a sample, but only generate high contrast intensity information via interference (Popescu et al. 2008a). Recently, there have been escalating interests in measuring 3-D RI distributions for various applications in biological imaging. Mainly because RI, as the primarily intrinsic optical parameters, provides the possibility for label-free live cell imaging with the capability of providing quantitative information about the sample. Although 3-D RI tomography does not provide molecular specificity in general, some specimen having distinct RI values such as lipid droplets (Kim et al. 2016b) or gold nanoparticles (Kim et al. 2018a; Sung et al. 2018) in the cytoplasm, can be specified and quantified.

Furthermore, 3-D RI tomography provides quantitative imaging capability; cellular dry mass or cytoplasmic concentration can be precisely quantified from the measured RI values, which are inaccessible with fluorescence imaging techniques. Most importantly, 3-D RI tomography does not require the use extracellular agent, and thus, it simplifies sample preparations and is also suitable for long-term live-cell imaging (Barty et al. 2000; Lee et al. 2013; Majeed et al. 2016; Popescu 2011). In addition, due to the quantitative and thus reproducible imaging capability, the use of 3-D RI tomography for the disease diagnosis is being actively investigated in combination with other techniques such as microfluidics (Merola et al. 2012; Sung et al. 2014), machine and deep learning algorithms (Jo et al. 2019; Jo et al. 2015; Jo et al. 2014; Jo et al. 2017; Rivenson et al. 2017), and fast imaging processing algorithms (Kim et al. 2013). In this mini-review, we introduce the principle of optical techniques that measure 3-D RI tomograms and summarize recent applications for the study of various biological and medical applications.

### The principles of biological imaging using refractive index as imaging contrast

The RI of material is obtained by measuring the interactions between light and matter. One of the well-known RI measuring techniques includes a refractometer that obtains the average RI value of a solution or a surface plasmonic sensor used to measure the surface RI of metal (Willets and Van Duyne 2007). These refractometer techniques are suitable for measuring a sample with the homogeneous distribution of RI values, such as a transparent solution. However, it is technically challenging to measure a sample with an inhomogeneous distribution of RI values, such as biological cells or tissues (Liu et al. 2016). This is because light refracts and reflects at the interface between the two media with different refractive indexes. When light refraction and reflection occurs many times, the coherence summation of these events can be expressed as multiple light scattering (Cheong et al. 1990). This explains why biological tissues appear opaque white, while individual cells seem transparent.

Conventionally, phase contrast or differential inference microscopy have been utilized to exploit RI distributions in samples. When a laser beam passes through a transparent specimen, such as individual biological cells, the laser beam acquires a distorted wavefront or phase information. This occurs because the speed of light passing through a specific part of the sample differs from another part due to inhomogeneous distribution of RI of the sample. Unfortunately, conventional image sensors do not directly measure this wavefront information, because the speed of light is much faster than the capturing ability of an image sensor. Thus, phase contrast or differential inference microscopy exploits the principle of light interference. Significant light interference can be created in an optical imaging system, such as bright-field microscopy, by inserting additional optical components(Lee and Park 2014). This allows conversion of wavefront information into intensity information that can be measured by an image sensor. This is how one can achieve high imaging contrast when imaging transparent biological cells using phase contrast or differential inference microscopy. Hence phase contrast or differential inference microscopy enables clear visualization of the boundaries of the cell membrane as well as subcellular organelles (Mann et al. 2005; Smith 1955; Zernike 1942; Zernike 1955).

However, these conventional interference microscopes, such as phase contrast or differential interference microscopy, can only provide qualitative information. This is because the relationship between wavefront information and intensity images for these interference microscopes is not straightforward, thus making it difficult to extract quantitative information(Smith 1955; Zernike 1942; Zernike 1955). Quantitative phase information can provide valuable information about the sample without using exogenous labeling agents. For example, the measurements of quantitative phase maps of red blood cells can be directly converted into a cell height information (Ikeda et al. 2005; Park et al. 2006; Popescu et al. 2006).

Various quantitative phase imaging (QPI) techniques have been developed and utilized for various research fields (Park et al. 2018d). In particular, Mach-Zehnder or Michelson types of interference microscopic techniques have been extensively utilized. In-line holography techniques simplify the optical setup by removing a reference arm. Quantitative phase microscopy techniques based on the transport of intensity or ptychography have provided enhanced imaging quality with relatively simple instrumentations. Recently, the QPI unit was developed as a filter-type add-on unit, which can be attached to convert a conventional bright-field microscope into a quantitative phase microscope (Lee and Park 2014). The detailed information on QPI and its application to biological studies can be found in elsewhere (Lee et al. 2013; Majeed et al. 2016; Popescu 2011).

### The principle of measuring 3-D RI tomography of cells and tissues

Even though 2-D QPI techniques provide quantitative and label-free imaging of live cells, they only provide topographic information; i.e., the measured optical phase delay is a coupled parameter of cell height and its RI distribution (Rappaz et al. 2005). Previously, several methods have been suggested in order to decouple the height and RI information in 2-D QPI techniques. Measuring two holographic images obtained with two different extracellular media with different RI values (Rappaz et al. 2005) or illumination with two wavelengths (Rappaz et al. 2008) provides the separation of height and RI from the measured holograms. Alternatively, the mean RI of suspended cells is calculated from measured 2-D optical phase delay images, assuming spherical shapes of cells (Kemper et al. 2007).

In order to measure 3-D RI tomograms of cells, various approaches have been demonstrated (Kim et al. 2016c). Among them, the angle scanning approach has been widely utilized (Fig. 1). First, multiple 2-D holograms of a sample are measured at various angles of illuminations [Figs. 1(a)-(b)], from which a 3-D RI tomogram of the sample can be reconstructed via inverse scattering theory. According to the scattering theory, the difference in wavevectors of the incident and scattered lights determine the spatial frequency information of the optical scattering potential of the sample in 3-D Fourier space [Fig. 1(c)]. Finally, a 3-D RI tomogram of the sample [Fig. 1(d)] is reconstructed by applying 3-D inverse Fourier transform. This technique has been widely known as optical diffraction tomography (ODT) and holotomography (HT). The principle of 3-D RI tomography is very similar to X-ray computed tomography (CT) where multiple 2-D X-ray images of the human body are measured at various illumination angles, and a 3-D X-ray absorptivity tomogram is then retrieved via the inverse scattering theory. Both X-ray CT and laser HT shares the same governing equation – Helmholtz equation, the wave equation for a monochromatic wavelength (Kim et al. 2016c).

**Figure 1.**
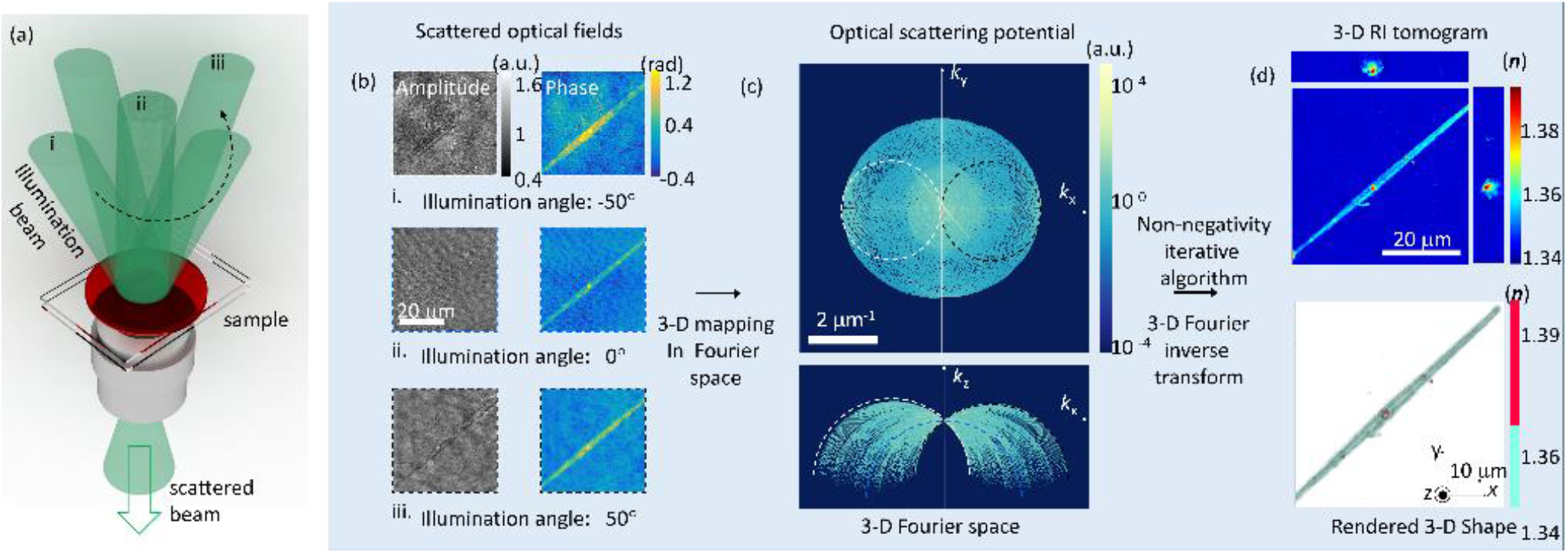
The schematic of 3-D refractive index tomography. (a) The illuminated plane waves and the scattered waves by the sample with distorted wavefronts. Three representative incident and scattered waves are depicted. (b) The retrieved 2-D complex optical fields of the light scattered by the sample. (c) The synthesized 3-D optical scattering potential in 3-D Fourier space. The regions corresponding to the three illumination angles are specified with distinct colors. (d) Three crosssectional slices and the iso-surface of the reconstructed 3-D refractive index tomogram.

The history of HT goes back to the late 60s. After the rise of X-ray CT technology and the invention of lasers, some pioneering physicists realized that the X-ray CT principle could also be applied to a laser. The principles of X-ray CT are based on wave propagation and can be described by the wave equation, except they use different wavelengths of waves. The first theoretical work was presented in 1969 (Wolf 1969), and the first experimental demonstration was shown in 1979 (Fercher et al. 1979). However, it seems that at the time many researchers did not realize this HT technique can be applied to biological imaging with significant benefits. The early applications of HT technique had been limited to measuring 3-D shapes of transparent microscopic plastics. In the 2000s, several research groups have revised and employed the HT technique for biological applications (Choi et al. 2007; Kim et al. 2014a; Lauer 2002).

HT technology directly provides the measurements of the 3-D RI distribution of a cell. The reconstruction of 3-D RI tomogram of the specimen is achieved by inversely solving the Helmholtz equation from a set of multiple 2-D optical field images of the sample. This set of 2-D optical field images of the sample containing the 3-D information is generally obtained by varying the illumination angle of a laser impinging onto the specimen (Choi et al. 2007; Kim et al. 2014c; Shin et al. 2015), or rotating the specimen while keeping the light source fixed (Barty et al. 2000; Charrière et al. 2006a; Kuś et al. 2014). For further information, the principle of HT, the detailed procedure with a MatLab code, the phase retrieval algorithms, and the regularization algorithms can be found elsewhere (Debnath and Park 2011; Kim et al. 2014a; Lim et al. 2015) summarize various regularization algorithms used in HT.

From a technical point of view, significant technical advancements have been made recently. For example, the sub-100-nm spatial resolution was achieved using the deconvolution of complex optical field (Cotte et al. 2013). Tomographic RI reconstruction with white light illumination was presented, which demonstrates a signification reduction of speckle noise and dramatically improved image quality (Kim et al. 2014b). Hyperspectral HT was also demonstrated; it measures 3-D RI tomograms of a sample at various wavelengths using a wavelength scanning illumination (Jung et al. 2016). The realtime reconstruction and visualization were also demonstrated, which was powered by a graphics processor unit (GPU) (Kim et al. 2013). It is worthy to note that in the mid-2010s, HT technology was commercialized and had started being used in biological laboratories and medical hospitals. As of 2018, two companies provides the commercialized HT systems – Nanolive (www.nanolive.ch) and Tomocube (www.tomocube.com).

### Opportunities and challenges of RI as imaging contrast

Exploiting RI as imaging contrasts have advantages and limitations. In this section, we summarize representative features of HT technology.

1. Label-free: Because RI is an intrinsic optical parameter of material. No labeling agents or dyes are required for imaging biological cells and tissues. It means 3-D images of live cells can be obtained for a long time as long as physiological conditions are met. In addition, it can save time and cost for sample preparation. This label-free feature might become powerful for some applications where cells are to be reinjected to human bodies, for example, as in immune therapy (Yoon et al. 2017; Yoon et al. 2015) or stem cell therapy (Braydich-Stolle et al. 2005).
2. Quantitative biological imaging: Using HT technology, RI value can be precisely measured. Unlike fluorescence techniques, where the intensity of fluorescence emission highly depends on protocol, and the results are only qualitative, HT technology provides highly reproducible RI value in a quantitative manner. Importantly, the RI value can be directly translated into the protein concentration information (Barer 1953). Furthermore, the dry mass of subcellular structures or cells can also be calculated from RI distributions (Barer 1953; Popescu et al. 2008b; Zangle and Teitell 2014).
3. Potentials in biological and medical applications: From spatial RI distributions, HT technology can retrieve various quantitative parameters. Certain molecules such as lipid(Jung et al. 2018; Kim et al. 2016b) or metal nanoparticles (Kim et al. 2018a; Turko et al. 2013) have distinctly high RI values, which can be addressed by measuring 3-D RI maps. RI values of a solution are linearly proportional to its concentration (Yoon et al. 2017). Local RI values in cells can be converted into cytoplasmic protein concentration (Barer 1952). The integration of RI values over a cell volume can be translated into dry cell mass. Furthermore, these RI values are intrinsic quantitative parameters of live cells, and thus can be utilized for a biophysical marker. For example, 2-D and 3-D RI distributional maps can be utilized in label-free cellular identification (Chalut et al. 2012; Jo et al. 2015; Jo et al. 2014; Jo et al. 2017; Yoon et al. 2017) and long-term growth monitoring (Bettenworth et al. 2014; Chalut et al. 2012). In addition, RI itself can be used as a new marker which represents cellular states or abnormalities (Lenz et al. 2013; Schürmann et al. 2016), and the factors which change the intracellular RI are being actively studied (Ekpenyong et al. 2013; Wang et al. 2011b).

HT technology also has several limitations and challenges:

1. Limited molecular specificity: Although RI values can be precisely measured using HT technology, it is difficult to relate these measured RI to molecular information. This is mainly because proteins have similar RI values regardless of their types. Nonetheless, the spatial distribution RI values can provide limited morphological information about subcellular organelles. For example, nucleus membrane, nucleoli, lipid droplets, and vacuoles can be easily identified from RI distributions because they have distinct RI values different from neighboring environments (Barer 1953; Jung et al. 2018; Kim et al. 2016b; Schürmann et al. 2016). Furthermore, the gradient of RI can also be utilized for further discriminating subcellular structures (Kim and Park 2018).

With these potentials and challenges, measuring RI tomography of various types of cells have become widely adapted for the study of various research topics. Below we highlight recent applications of measuring RI tomograms for various research disciplines.

## Microbiology

Observing individual bacteria with conventional optical microscopes is challenging. This is due to a number of reasons. One is the size of most bacteria, which are in the order of micrometers or smaller. In addition, bacterial cytoplasm has refractive indexes similar to that of a medium, and the bacteria are transparent. Therefore it is difficult to visualize bacteria under a conventional bright field microscope and a high-resolution microscope is required.

When imaging a bacterial cell using a phase contrast microscope, it is possible to obtain good imaging contrast. In this case, however, only limited morphological information is provided, such as the length, width, and shape of bacterial cells. Although phase contrast microscopy can measure live bacteria without using labels, it has limited capability for the study of microbiology because it provides only 2-D qualitative images. Immunofluorescent labeling technology can provide molecular-specific information inside bacteria, but it can be time-consuming and costly to stain the cells. In addition, research is limited due to secondary problems such as photochromism and phototoxicity that occur during the bleaching process. This makes it difficult to observe the bacteria for long periods of time. Traditionally, confocal microscopes and transmission electron microscopes have been used to obtain the internal structure of individual bacteria at high resolution. However, it is difficult to observe live cells for a long time because these techniques require cell staining or fixation.

RI tomographic imaging techniques can solve the problems of these existing imaging techniques. Because it is a non-invasive, label-free method, living bacteria can be observed for a long time without using additional exogenous labeling agents. In particular, by measuring the RI, protein concentration and mass information within the bacteria can be extracted, which has recently led to several studies related to the division of bacterial cells. However, because RI itself does not provide molecular specific information, an in-depth investigation in the context of molecular biology is significantly limited. In the future, there will be potentials where both 3-D RI tomographic imaging and fluorescence microscope technique are used simultaneously.

Several previous papers had been reported where 3-D RI tomograms of individual bacteria are measured using HT technology. Using Mach–Zehnder interferometer and illumination scanning, 3-D RI tomograms of bacteria extracted from a sample of stool (Lauer 2002) and *E. coli* (Cotte et al. 2013) have been reported. Recently, white-light diffraction tomography was used to image 3-D RI tomogram of *E. coli* (Kim et al. 2014b). More recently, 3-D RI distribution of *Magnetospirillum gryphiswaldense,* a magnetotactic bacterium which produces magnetic particles (magnetosome), were measured (Bennet et al. 2016).

RI information can be exploited to retrieve cellular dry mass and concentration information about individual bacteria. Dry mass refers to the non-aqueous contents inside cells and can be used as an indicator of cellular growth and division. Because RI of cytoplasm is linearly proportional to its concentration, RI tomography represents protein distribution of a cell. Furthermore, the integration of RI over cell volume can also provide information about the dry mass of the cell. Dry mass of a cell can be simply retrieved by measuring 2-D optical phase delay maps and averaging it over cell area because the optical phase delay map of a cell corresponds to the integration of RI differences between non-aqueous contents inside cells (Lee et al. 2013; Popescu et al. 2008b). The cellular dry mass of fission yeast was monitored during the cell cycle with digital holographic microscopy, and the difference of mass production rate between wild-type and mutant fission yeast cells was observed (Rappaz et al. 2009). Using spatial light interference microscopy (SLIM), the dry mass of *Escherichia coli* cells was measured (Mir et al. 2011). In this work, the roles of cell density and morphology in mass regulation had been investigated.

The application of RI based imaging in microbiology especially has a strong advantage for the long-term growth monitoring of microbes because RI provides both the morphological information and quantitative information about cell mass without using exogenous labels. Recent reports show the bacterial species identification using the 2-D optical phase delay maps and artificial intelligence algorithms (Jo et al. 2015; Jo et al. 2014; Jo et al. 2017). More recently, 3D RI tomograms were utilized for evaluating antibacterial activities of the graphene-based film (Kim et al. 2017b).

### Hematology

The physical parameters of red blood cells (RBCs) are strongly related to the pathophysiology of various diseases (Suresh 2006). Conventionally, optical microscopy with labeling methods has been used to examine RBC morphology in blood smears; poikilocytosis (e.g., spherocytes, target cells) and blood-borne infectious diseases including malaria are routinely examined. Information about hemoglobin (Hb) in RBCs are of particular importance in laboratory medicine; mean corpuscular Hb concentration (MCHC) and mean corpuscular Hb content (MCH) are extensively examined for medical diagnosis. It is also well known that the deformability of RBCs can be altered by several infectious diseases and genetic disorders (e.g. malaria and sickle cell disease) (Byun et al. 2012; Diez-Silva et al. 2012; Kim et al. 2014d; Mills et al. 2007; Park et al. 2008), with the implications of malfunctions in microcirculation. In clinical hematology, automated blood cell counters based on the complete blood count (CBC) have been utilized to measure the properties of RBCs. Current automated blood cell counter techniques measure the parameters of RBCs, including mean corpuscular volume (MCV), MCHC, MCH, and RBC distribution width (RDW), which serve as the principal and crucial information from which clinicians diagnose abnormalities in RBCs.

The use of 3-D RI tomography in the field of hematology could lead to the simultaneous measurements of various optical parameters of individual RBCs. Figure 2 summarizes the analysis procedure for retrieving the parameters of individual RBCs using 3-D RI tomography, including the volume, surface area, sphericity, Hb content, Hb concentration, and membrane fluctuation, which can be obtained at the single cell level. Figure 2 summarizes the analytical procedure for retrieving the parameters of individual RBCs using 3-D RI tomography, including volume, surface area, sphericity, Hb concentration, Hb content, and membrane fluctuations.

**Figure 2.**
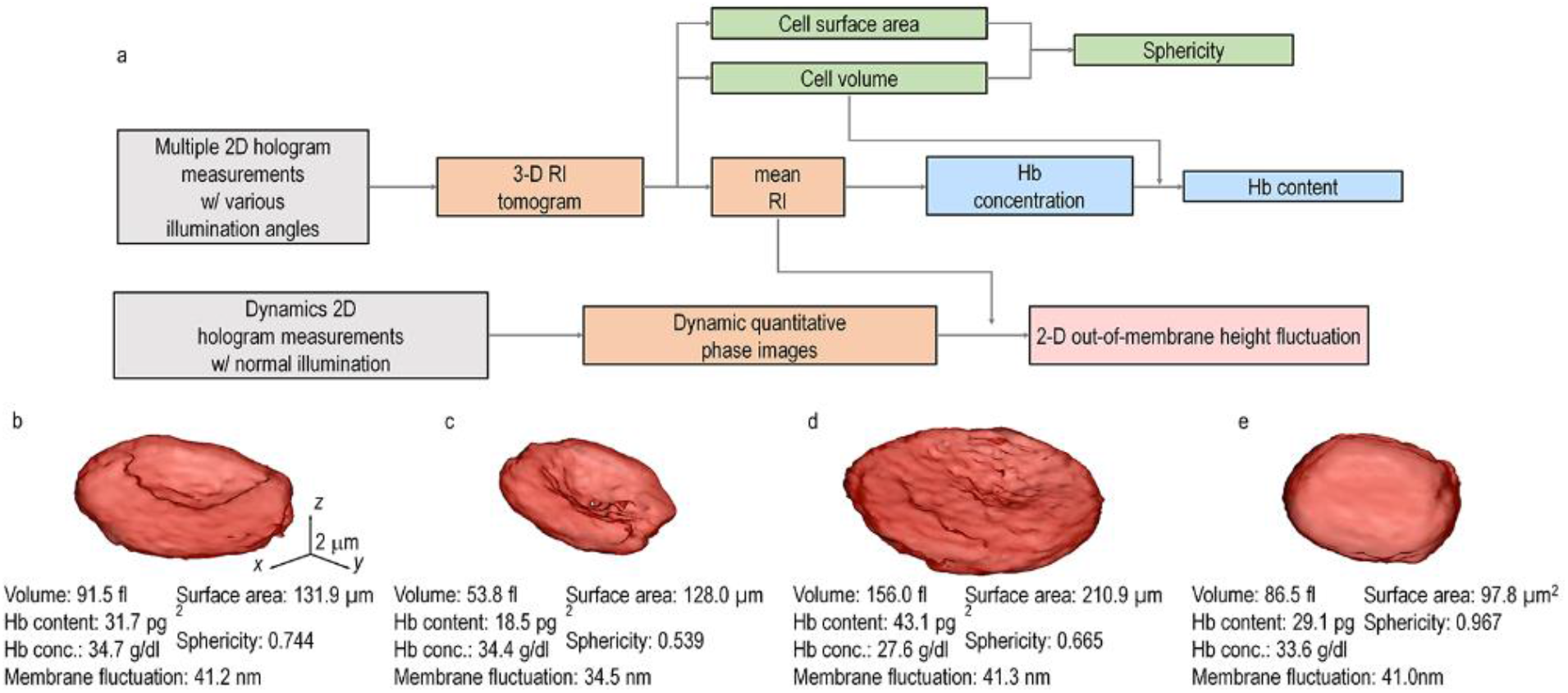
(a), Schematic diagram of the analysis procedure for retrieving the parameters of individual RBCs using 3-D RI tomography. (b-d), 3-D rendered isosurfaces of RI tomograms of individual RBCs and retrieved parameters from (b) healthy, (c) Iron-deficiency anemia, (d) reticulocyte, and (E) HS red blood cells. Reproduced from Ref. (Kim et al. 2014d) with permission.

The measurements of various parameters of RBCs enables single-cell profiling, of which the importance has escalated in recent years (Higgins 2015; Weatherall 2011). Regulation of dynamic cellular systems and related pathophysiology can be better understood when the various parameters of individual cells are simultaneously analyzed in detail. This is because of recent improvements in measurement techniques, which have made single-cell profiling more efficient than before. In addition, data-based research approaches in biology and medicine have also grown rapidly, allowing researchers to explore new perspectives, in addition to existing hypothesis-based research methods. The 3-D RI measurements enable the retrieval of morphological (cell volume, cell surface area, sphericity), biochemical (Hb concent, Hb concentration), and biomechanical (dynamic membrane fluctuation) information measured at the individual cell level. This allows correlative analysis, which is not possible with the conventional blood cell counters.

Previously, 2-D QPI techniques have been employed for various applications in hematology. In particular, optical measurements of the parameters of individual RBCs have been widely studied, including malaria-infected human red blood cells (Chandramohanadas et al. 2011; Park et al. 2008), sickle cell diseases (Byun et al. 2012; Shaked 2012), and ATP-dependent fluctuations (Park et al. 2010). RBCs do not have subcellular organelles and exhibit homogeneous RI distributions, which therefore allows measurements of Hb concentration. Thus, 2-D QPI measurements can also provide both the morphological information (cell height map) as well as the biochemical information (Hb contents or dry mass), with prior information about Hb concentration or the RI value of RBC cytoplasm. However, RBCs from an individual with diseases such as malaria infection or sickle cell anemia, the values of Hb concentrations, and thus RI values, vary significantly. Therefore, it is required to directly measure the 3-D RI tomograms of individual RBCs for the systematic study of disease states.

Recently, 3-D RI tomograms of RBCs have been utilized for measuring the parameters of individual RBCs. Lee et al. used common-path diffraction optical tomography (cDOT) (Kim et al. 2014c), and measured 3-D RI tomograms and dynamic membrane fluctuations of RBCs exposed to ethanol (Lee et al. 2015). It was observed that RBCs exposed to an ethanol concentration of 0.1–0.3% v/v becomes more spherical shapes than those of normal cells (Lee et al. 2015). Using the cDOT, the properties of individual RBCs stored with and without a preservation solution, citrate phosphate dextrose adenine-1(CPDA-1) were reported. In this work, various red blood parameters were analyzed and the results showed that, in the absence of CPDA-1, RBCs undergo a dramatic morphological transformation from discocytes to spherocytes within two weeks (Park et al. 2016). The RBCs also experienced a reduction in cell surface areas and became less flexible. However, RBCs stored with CPDA-1 retained their biconcave shapes and deformability for up to six weeks.

More recently, the RBCs from patients with diabetes mellitus have been systematically measured using 3-D RI tomography and dynamic membrane fluctuations (Lee et al. 2017). Morphologies of the RBCs from diabetic patients were not significantly different from the healthy ones. The deformability of the RBCs from diabetic patients was significantly lower than those of healthy RBCs, which is consistent with the previous literature using ektacytometry or filtration assay. Interestingly, this work reported the negative correlation between dynamic membrane fluctuation and glycated Hb concertation or HbA1c level, for the healthy RBCs; the higher the HbA1c level, the less deformable the cells are. In addition, the alterations in RBCs resulted from the binding of melittin, the active molecule of apitoxin or bee venom, have been studied by measuring 3-D RI maps and dynamic membrane fluctuations (Hur et al. 2017). RBCs from the cord blood of newborn infants and adult mothers or nonpregnant women were also systematically studied using 3-D RI tomography (Shin et al. 2015).

The study of white blood cells using 3-D RI tomography has not been fully exploited, but it will open new applications. More recently, Yoon et al. measured 3-D RI tomograms of mouse lymphocytes and macrophage (Yoon et al. 2015). In this work, the morphological alternations in lymphocytes, caused by lipopolysaccharide which is known to immunologically stimulate lymphocytes, were analyzed.

The applications of 2-D and 3-D RI imaging techniques in hematology has increased. Among them, measurements of membrane fluctuation dynamics and Hb concentration of individual RBCs have been actively investigated due to their correlations with diseases and environmental abnormalities. In addition, because various cellular parameters are simultaneously extracted from individual blood cells, and their correlative analysis can be followed, more previous investigation of cellular alterations associated with various diseases are now accessible with direct experimental approaches.

### Infectious diseases

The visualizations of the structures and dynamism of parasites and host cells are important for the study of parasitic infections. Electron microscopy provides spatial resolutions much higher than optical microscopy, and has been used to visualize the internal structures of parasites. However, the high spatial resolution of electron microscopy comes at the cost of static imaging; it does not provide the time-lapse information of parasitic infections. Optical microscopy techniques have been extensively used for imaging parasites and host cells (Cho et al. 2011). The use of labeling agents with high molecular specificity have elucidated the molecular biology of various infectious diseases. However, conventional fluorescent labeling techniques only provide qualitative imaging capability, and some parasite is difficult to be labeled.

Recently, 3-D RI tomography techniques have been utilized for the field of infectious diseases. Park et al. have used tomographic phase microscopy and measured 3-D RI maps and dynamic membrane fluctuations of malaria-infected RBCs as a function of various infection stages (Park et al. 2008). *Plasmodium falciparum* parasites invaded into host human red blood cells were visualized from the measurements of 3-D RI tomography. Also, the significantly decreased dynamic membrane fluctuations in the infected RBCs were also reported, indicating the decreased cell deformability (Diez-Silva et al. 2010; Diez-Silva et al. 2012). The label-free capability of 3-D RI tomography has been utilized for the study of egress of malaria parasites (Chandramohanadas et al. 2011), which provided a comprehensive body of information on the relationships between biomechanical and biochemical parameters and parasite egress from RBCs infected by malaria-inducing parasites.

Kim et al. employed the ODT algorithm to reconstruct 3-D RI tomography of malaria-infected RBCs and it shows better image quality compared to the ones obtained with optical projection algorithm because ODT considers light diffraction inside samples(Kim et al. 2014a). In addition, various morphological information about invading parasites and produced hemozoin structures are obtained and analyzed quantitatively. The evasion mechanisms of *P. falciparum* from host immunity was also studied using holotomography (Tougan et al. 2018).

More recently, *Babesia microti* invaded RBCs were investigated by measuring 3-D RI tomograms at the individual cell level (Park et al. 2015). *B. microti* causes emergency human babesiosis, which shares similar pathophysiology and pathologic symptoms with malaria. In this work, RI information was effectively used for the study of babesiosis, because the RI of *B. microti* parasites is distinct from RBC cytoplasm can thus be clearly visualized, otherwise very difficult to be identified with conventional optical imaging techniques because of the lack of effective labeling agents for *B. microti* parasites. Ekpenyong et al. reported the RI maps of Primary murine bone marrow-derived macrophages which were infected by *Salmonella enterica* serovar Typhimurium (Ekpenyong et al. 2013).

3-D RI tomography was also employed for the study of viral infection. Simon et al. used a setup, in which fluorescence confocal microscopy and optical diffraction tomography were combined, and studied human respiratory epithelial carcinoma A549 cells infected with human influenza H3N2 virus (Simon et al. 2010). Interestingly, in the infected cells, the spherical structures with the size of 150–200 nm and with the distinctly high RI values were observed, which were expected to correspond to the buddings of viral particles.

In the field of microbiology, various cases have been studied where parasites can be detected using their distinctive RI values. Also, alterations in infected host cells are quantitative studies by measuring 3-D RI distributions. RI-based imaging techniques can potentially provide advantages over conventional techniques such as chemical assay and fluorescence microscopy, because of its label-free imaging capability and simple sample preparation procedures.

### Hepatology

Optical microscopic imaging of hepatocytes has played an important role in hepatology. The structures of cells and sub-cellular organelles and their dynamics are strongly correlated to the physiology of hepatocytes, and also significantly altered associated with liver-related diseases. Recently, Kim et al. have measured 3-D RI tomograms of human hepatocytes (human hepatocellular carcinoma cell line, Huh-7) were measured at the individual cell level (Fig. 3). In this work, various subcellular structures of hepatocytes are clearly identified using RI values, including cell membrane, nucleus membrane, nucleoli, and lipid droplets (LDs) (Kim et al. 2016b). Also, time-lapse 3-D RI tomograms of hepatocytes were also measured, from which dynamics of individual LDs were quantified.

**Figure 3.**
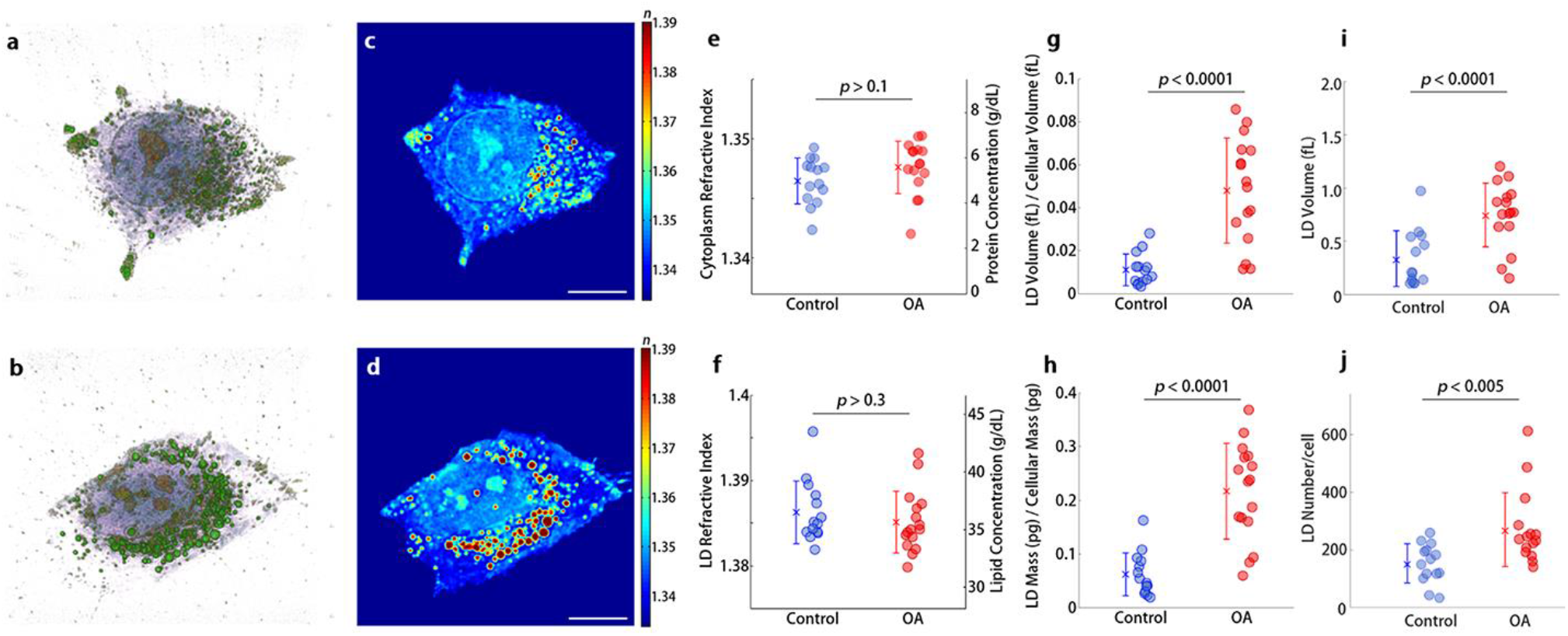
**(a,b)** 3-D rendered isosurface image of 3-D RI distribution of (**a**) an untreated and (**b**) oleic acid (OA) treated hepatocyte. (**c,d**) Cross-sectional slice images of 3-D RI distribution of (**c**) the untreated and (**d**) OA treated hepatocyte. The scale bars indicate 10 μm. e-j, Quantitative analysis of (**e**) RI of cytoplasm, (**f**) RI of LDs, (**g**) the ratio of LD volume to cell volume, (**h**) the ratio of LD mass to cellular dry mass, (**i**) volume of individual LDs, and (**j**) number of LDs in untreated and OA-treated hepatocytes. Reproduced from Ref. (Kim et al. 2016b) with permission

Among subcellular structures of hepatocytes, lipid droplets (LDs) are of particular interest because they are directly related to the lipid metabolism. LDs consist of a monolayer of phospholipids and associated proteins surrounding a core of neutral lipid and are ubiquitous intracellular organelles storing and supplying lipids in most cell types as well as hepatocytes

(Martin and Parton 2006). Recent studies suggest that LDs participate in various pathological roles, such as cancer and diabetes mellitus, and exhibit 3-D motions to regulate lipid storage and metabolism (Welte 2009). However, the detailed process of LDs dynamics including biogenesis, growth and 3-D subcellular motions are incomplete (Wilfling et al. 2014).

LDs can be effectively visualized exploiting its RI value; the RIs of lipid are significantly higher than protein(Beuthan et al. 1996) and thus LDs can be identified by measuring 3-D RI tomograms. Measuring LDs using the high RI values have several advantages over conventional approaches. Fluorescence techniques for labeling LDs have been widely used (Martin et al. 2005), but the use of fluorescent probes requires for ethanol treatment, which can influence the physiological conditions of LDs such as induced fusions of LDs in live cells (Fukumoto and Fujimoto 2002). Coherent anti-Stokes Raman scattering Raman scattering techniques have been used to visualize LDs in live cells without the use of exogenous labels (Evans et al. 2005; Nan et al. 2003), but it requires for highly expensive laser and detection instruments and have the issue of low signal-to-noise ratio for molecules with low concentrations. The recent study demonstrated that the time-lapse 3-D RI distribution of LDs in live hepatocytes could be quantitatively analyzed with ODT (Kim et al. 2016b). The shapes, sizes, and the masses of individual LDs in live hepatocytes were retrieved from the measured 3-D RI tomograms.

One of the direct and important applications of HT technology would be imaging and quantifying individual lipid droplets in hepatocytes. 3-D RI distributions of hepatocytes can provide quantitative imaging and analysis about metabolisms of lipids droplets in live cells. In addition, 3-D RI maps of live cells also provide morphological information, and thus the generations, dynamics, and degradation of lipid droplets can also be studied with the combination with the information about subcellular structures.

### Histopathology

RI can potentially serve as an important contrast in histopathology because (i) the use of RI for imaging tissue slides provide imaging contrast for the visualization of anatomical features in tissue slides otherwise invisible under conventional bright-field microscopy and (ii) it does not require for the labeling process which can save time and cost. In addition, unlike conventional histopathology that relies on staining agents, the use of RI can provide quantitative criteria for pathologies in unlabeled biological tissues.

The RI information, for example, was used to quantify the weight loss in the inflammation-induced colitis (Lenz et al. 2013). Besides the dry mass information, the measurements of optical phase delay maps of a tissue slice could precisely be utilized to extract the scattering parameters such as scattering coefficients (*μ_s_*) and anisotropies (*g*) (Ding et al. 2011; Wang et al. 2011a; Wang et al. 2011b). This is because light scattering in tissue is caused by inhomogeneous distributions of RI, and thus as light passes through tissue, it undergoes significantly large events of light refraction and reflection, resulting in complex patterns of multiple light scattering. These scattering parameters have been employed to investigate the morphological alterations in prostate cancers (Figs. 4a-b) (Sridharan et al. 2014), breast cancers (Majeed et al. 2015) and epithelial pre-cancers(Su et al. 2015), which can be potentially utilized for diagnostic purposes.

**Figure 4.**
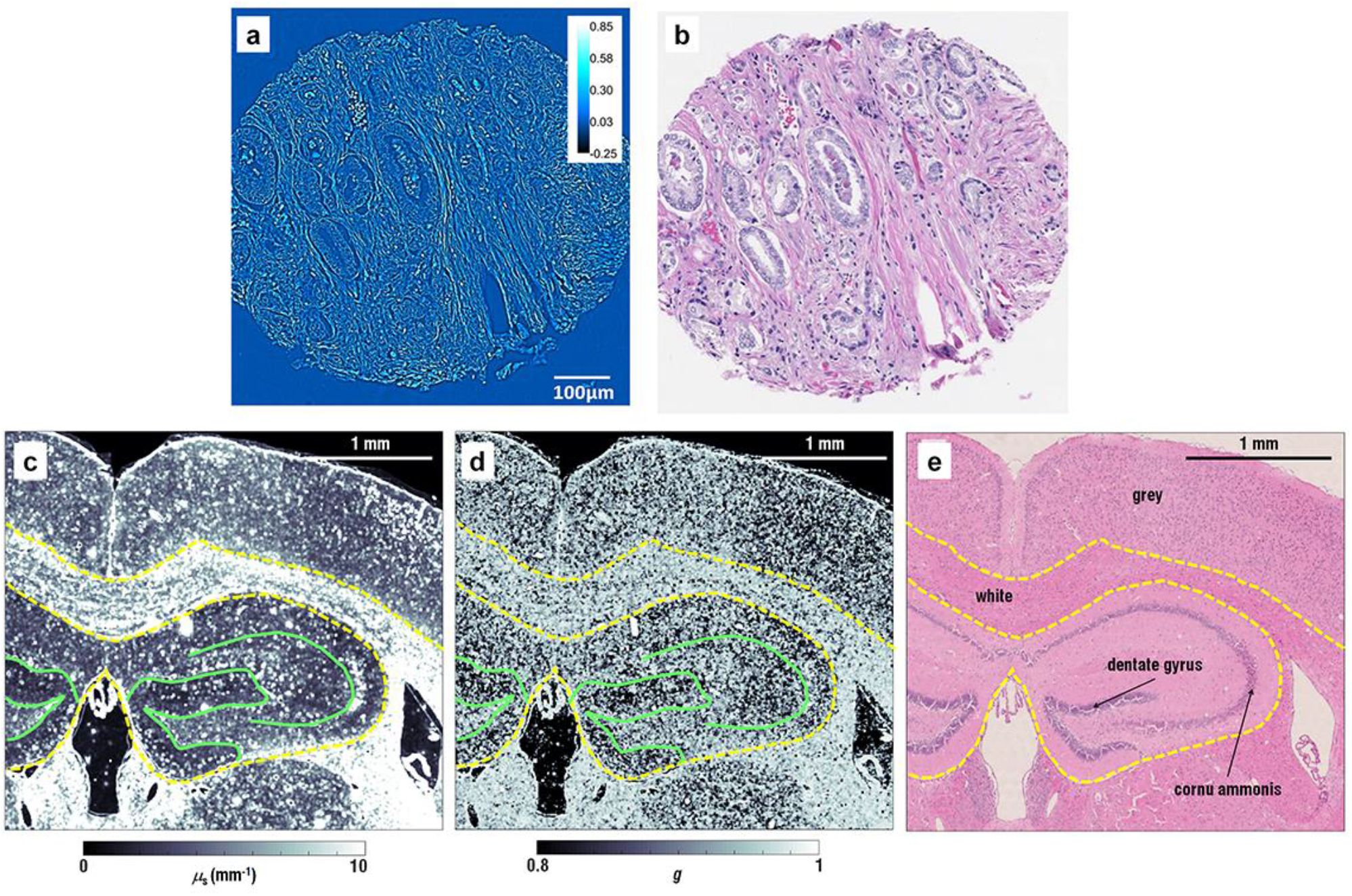
Applications of RI-based imaging techniques for histopathology and cell dynamics (a) 2-D optical phase map of a slice from a biopsy sample obtained from a patient who had a biochemical recurrence of prostate cancer after undergoing radical prostatectomy. (b) The adjacent tissue slice of (a) labeled with conventional H&E staining. (c-d) Scattering parameter (*μs*) map (c) and anisotropy (*g*) map (d) of a whole mouse brain tissue slice, which was retrieved from the measured 2-D optical phase map of the slice. (e) The adjacent tissue slice labeled with conventional H&E staining. Reproduced from Refs. (Lee et al. 2016) and (Sridharan et al. 2014) with permission.

Recently, the label-free tissue imaging capability by utilizing RI information was adapted to neuroscience. Optical phase delay maps of brain tissues were obtained for whole brain mouse tissue slides (Lee et al. 2016). The maps of scattering parameter (*μ_s_* and *g*), extracted from the measured optical phase delay images of brain tissue, showed anatomical structures, comparable to the ones obtained with the conventional hematoxylin and eosin (H&E) staining method (Figs. 4c-e). Furthermore, these scattering parameter maps showed a statistical difference between the tissue slides from mice with Alzheimer’s disease and the ones from healthy mice, indicating the structural alternations in the tissue.

For the application in histopathology, RI-based imaging of tissues can provide various parameters such dry mass, scattering coefficients, and anisotropies, and these values can be quantitatively analyzed in order to investigate morphological or structural alterations associated with diseases. In the near future, the use of QPI techniques will be further broadened for the diagnosis of various diseases (Kim et al. 2016a). Also, the RI-based imaging techniques does not require for complex procedures for sample preparation, which can dramatically reduce the time and cost for histopathological diagnosis (Greenbaum et al. 2014).

### Outlook

Here we presented the principles of quantitative phase imaging techniques which exploit RI as an intrinsic optical imaging contrast for cells and tissues. Although this is a relatively new research field, and only a few topics of the studies have been investigated so far, the research work reviewed here suggest that the use of RI for biological imaging may play an important role in various fields of studies (summarized in Table 1), where label-free and quantitative live cell imaging capability provides benefits. This would significantly enhance our understanding of the pathophysiology of various diseases, which may also lead to the development of novel diagnostic strategies in the future.

**Table 1.**
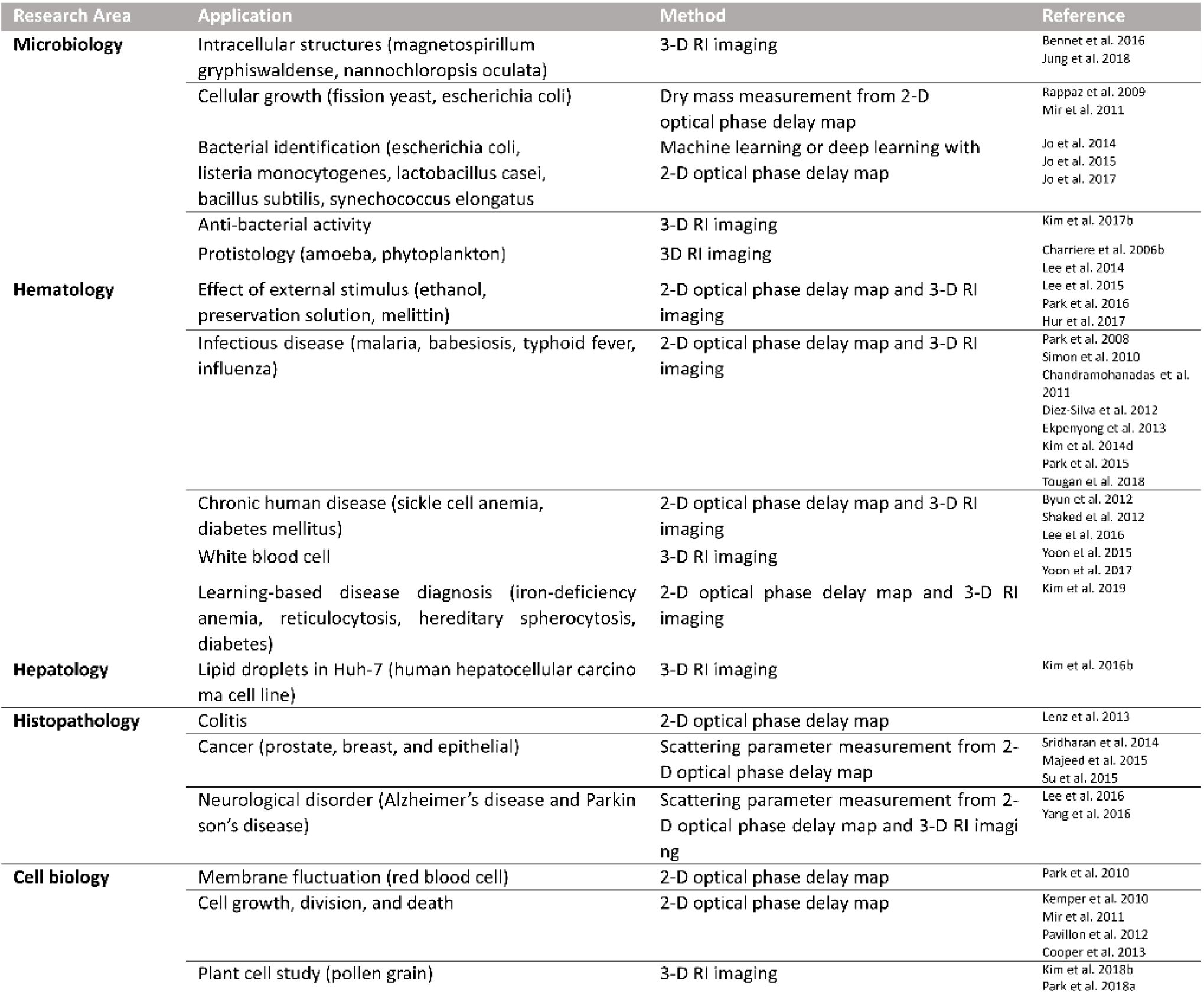
Major research areas which 2-D and 3-D RI measurement techniques have been employed.

The uses of RI as intrinsic imaging contrast for cells and tissues for biological and medical applications have not yet been fully explored; there are still various important issues in the physiology and pathology, which can be addressed by the utilization of RI-based imaging techniques and corresponding analysis methods. Some of the representative cell images in emerging fields are presented in Fig. 5. Many interesting studies which had been performed with 2-D QPI techniques would be readily investigated with 3-D RI tomography techniques because 3-D RI measurements directly provide both RI values and shape information, whereas usual 2-D QPI techniques only provide optical phase maps – a coupled parameter between RI values and the height of a sample. For example, water flux in individual neuron cells (Jourdain et al. 2011), morphologies of tumor cells (Kemper et al. 2006), cell growth and division (Cooper et al. 2013; Kemper et al. 2010; Mir et al. 2011), and cell death (Pavillon et al. 2012) would be studied in more details with 3-D RI tomography.

**Figure 5.**
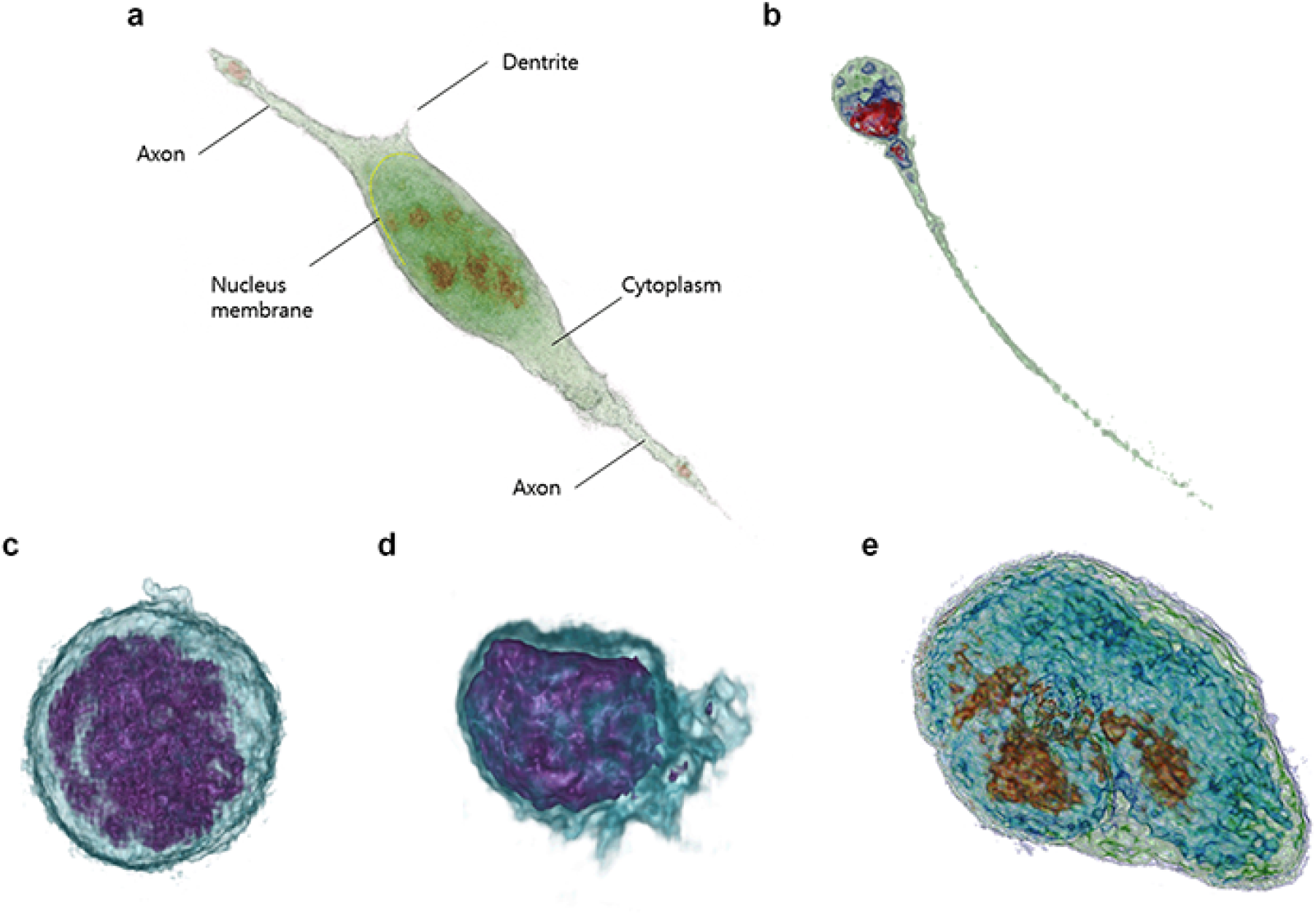
Emerging fields of research using 3-D RI tomography and representative cell images: (a) neuron cell; (b) human sperm; (c) mouse macrophage; (d) LPS-treated mouse macrophage; (e) embryonic stem cells. Image in (a) is reproduced from Ref. (Yang et al. 2016) with permission. Images in (b-e) are provided by Tomocube Inc.

Label-free live-cell imaging capability of 3-D RI tomography will be a strong advantage for its future applications in medicine and biology. For example, it has a potential to be used in the studies of stem and neuron cells which are vulnerable to environmental changes such as the use of fluorescent labeling agents (Braydich-Stolle et al. 2005; Millet et al. 2007). Although 3-D RI tomography does not generally have molecular specificity, it can provide information about the cellular morphology, internal structural changes, and various biophysical parameters. Thus, 3-D RI tomography will be a strong complementary technique to fluorescence techniques. For example, a multimodal approach of using both the 3-D RI tomography and 3-D fluorescence microscopy open new avenues for the study of cell biology (Kim et al. 2017a; Kim et al. 2018c). Recently, the combination of 3-D RI tomography and structured illumination microscopy for 3-D fluorescence imaging was demonstrated (Chowdhury et al. 2017; Shin et al. 2018). The specific molecules or proteins can be localized in live cells using 3-D fluorescence microscopy, and then long-term imaging of cell dynamics can be followed. Also, by utilizing 3-D fluorescence images, analyzing quantitative information about subcellular organelles such as concentration or dry mass can be performed. In addition, RI-based imaging has shown promises toward label-free cell identification. Although conventional fluorescence-based cytometry or imaging provides extremely high molecular specificity and signal-to-background ratio, they only provide qualitative information which is highly dependent of protocols, sample conditions, etc. In contrast, RI is an intrinsic optical parameter of material, and thus can serve as a highly reproducible and yet quantitative imaging contrast. When cleverly combined with appropriate algorithms, the 3-D or time-lapsed 3-D RI data obtained from individual cells can be effectively analyzed.

Recently, RI-based imaging data has begun to be exploited in combination with machine learning algorithms (Jo et al. 2019). For example, from the 2-D quantitative phase images of individual bacteria, the genus of various bacteria was distinguished using a machine learning algorithm (Jo et al. 2015; Jo et al. 2014). In addition, non-activated lymphocytes were recently identified and classified from their 3-D RI tomograms (Yoon et al. 2017). Recently, the weapons-grade anthrax spores are optically detected using 2-D QPI techniques and deep learning (Jo et al. 2017). Label-free classification of kinetic cell states was also demonstrated using 2-D QPI and machine learning (Hejna et al. 2017). More recently, parameters of 3-D QPI were exploited for the diagnosis of hematologic diseases with high accuracy (Kim et al. 2019). The combination of RI-based imaging techniques and machine learning will potentially generate strongly synergetic effects for the study of stem cells and neuroscience where the use of exogenous labeling agents are generally avoided (Maxmen 2017).

Finally yet importantly potential topics in the application would include the 3-D label-free imaging for the study of Protists where 3-D morphometry and spatial movement are important issues. Previously, 3-D RI tomography has shown potentials for the in-depth investigation of amoeba (Charrière et al. 2006b) and phytoplankton (Lee et al. 2014). Also, the imaging of plant cells could be an important topic of study where label-free characterization and quantification of subcellular structures provide useful information (Kim et al. 2018b; Park et al. 2018a).

To facilitate the application of RI-based imaging methods for toxicology and pharmaceutical industry, it is required to achieve a platform to obtain high-throughput data. Toward this direction, several research activities are currently ongoing. Kim et al., presented a method to significantly enhance the speed of 3-D RI tomography employing sparse illumination patterns and a graphics processor unit (GPU) (Kim et al. 2013). Sung et al. demonstrated a method to obtain 3-D RI tomography of cells flowing in a microfluidic channel; without rotating a sample nor illuminating different patterns, optical information to construct a 3-D RI tomogram is acquired from the translations of a sample in the channel (Sung et al. 2014). Also, optical manipulation and 3-D tracking of biological cells in microfluidic channels may also open new applications combined with 3-D RI tomography (Merola et al. 2012).

Developments in new analysis algorithm are also required. Previously, the quantitative imaging capability of RI-based imaging had been limited to static analysis, including cellular dry mass, morphological, and biochemical information. The RI information can also be exploited in analyzing the dynamics of cells and subcellular compartments. Measurements of time-lapse 3-D RI tomograms of cells would accompany with the developments in analysis algorithm (Ma et al. 2016).

New techniques to effectively handle the large size of data are particularly needed to make RI-based imaging approaches even more useful. In order to perform analysis at the individual cellular level and reveal the details of the underlying mechanisms of diseases, at least dozens of cells should be measured in each experimental group. Then a technical issue arises – the sizes of tomogram files are enormously significant. For example, HT-1 series from Tomocube Inc., a commercialized 3-D holographic microscopy system, generate a data file with a 67 Mega voxels for one 3-D RI tomogram; the imaging volume is 84.3 × 84.3 × 41.4 μm with the pixel resolution of 112 × 112 × 356 nm for x-, y-, and z-direction, respectively. With the single-precision (or single) data type, a single tomogram of 67 Mega voxels will have the size of 268 Mbyte. When a 4-D or time-lapse 3-D tomograms the frame number of 512 is recorded, the data size will be approximately 134 Gbyte. More advanced methods to transfer, handle, and store such large sized data will also be required in the future.

To provide better molecular sensitivity to RI-based imaging, multispectral approaches have been introduced recently (Jung et al. 2014). The use of multiple coherent lasers (Jang et al. 2012; Park et al. 2009) or wavelength-scanning illumination (Jung et al. 2013; Rinehart et al. 2012) enabled measuring the optical phase maps as a function of wavelength, and this information was utilized for molecular imaging because certain molecules have distinct optical dispersion properties, i.e., different RI values as a function of wavelengths. Recently, 3-D RI tomography has been achieved for a large number of wavelengths using a super continuum source and a wavelength-scanning unit (Jung et al. 2016). Besides the technical difficulty, the major challenge in exploiting optical dispersion for molecular specificity is that most molecules present in cells do not exhibit strong optical dispersion except specific molecules such as Hb (Jung et al. 2014). Alternatively, the use of an ultraviolet light source was presented to better visualize chromosomes inside a cell (Sung et al. 2012). More recently, the use of gold nanoparticles (GNPs) has been utilized in RI-based imaging, because the strong light absorption and scattering of GNPs at a resonant wavelength enable high imaging contrast (Kim et al. 2018a; Turko et al. 2013).

Besides the developments in optical instrumentations and correlative imaging strategy, the advancements in reconstruction algorithm are expected to further enhance the quality of 3-D RI tomography. The current reconstruction algorithms are mostly based on the first order approximations – the algorithm only considers single scattered light to inversely solve the Wave equations. This assumption is valid for imaging single or up to a few layers of biological cells. However, the reconstruction may fail for highly scattering samples, such as thick biological tissues or embryos. Recently, several methods have been introduced to consider high order scattering in the reconstruction algorithm, including a nonlinear propagation model based on the beam propagation method (Kamilov et al. 2016) and a treatment of an unknown surrounding medium as a self-resonator (Lim et al. 2017). Furthermore, the resolution of the optical imaging system measuring 3-D RI tomography is governed by the numerical aperture of the objective and condenser lenses (Park et al. 2018b). Due to the limited numerical apertures of an imaging system, a fraction of side scattering signals is not collected, resulting into a reduced spatial resolution and inaccurate values of RIs in a reconstructed tomogram – known as a *missing cone* problem. To remedy this issue, several regularization algorithms have been used based on prior information about a sample (Lim et al. 2015). However, the careful use of appropriate prior condition is required and the computing power for the existing regularization algorithms are heavy. The further developments are also expected to more effectively address this missing cone problem.

In addition, optical trapping techniques can be combined with RI-based imaging techniques to revolutionize the way in which biologists approach questions in the field of cell-to-cell interaction and mechanobiology. Toward to this direction, the combination with 3-D RI tomography and optical manipulation techniques are beneficial. The optical tweezer technique has been shown to optically trap spherical particles, which aided to manipulate individual cells. Recently, 3-D RI tomography technique was combined with holographic optical tweezers, which demonstrates the 3-D dynamic imaging of the interaction of an optically trapped particle to a macrophage (Kim et al. 2015). More recently, the 3-D RI maps of biological cells were measured in order to actively control optical wavefront for trapping beams, and the demonstration of stable control of complex shapes objects such as RBCs and dimers was demonstrated (Kim and Park 2017).

From an application point of view, the utilization of microfluidic devices for 3-D RI tomography may open various opportunities. The use of a designed chip can be used to simplify an optical setup for the 3-D RI measurements (Bianco et al. 2017), and to allow the quantification of chemical concentrations of fluids in a microfluidic channel (Park et al. 2017a). The label-free imaging using RI will also have potentials for the cytotoxicity assay (Kwon et al. 2018).

One of the future directions of RI-based imaging would be toward *in vivo* application. If the use of RI information can be collected *in vivo,* it would bring significant impacts to the early diagnosis of various diseases. However, because of multiple light scattering in biological tissues deteriorate the delivery of optical information (Yu et al. 2015), the direct application of a simple optical imaging system would not be able to acquire clear images. Recently, 3-D RI tomograms of individual RBCs flowing through microcapillaries (Kim et al. 2016a). However, in this work, the part of a thin mesentery tissue should be placed in 3-D holographic microscopy. Ford et al. demonstrated *in vivo* phase imaging by collecting *en face* phase gradient images of thick scattering samples (Ford et al. 2012). More recently, Laforest et al. demonstrated *in vivo* phase imaging of retinal cells using transcleral illumination, which gives a dark-field configuration with a high numerical aperture (Laforest et al. 2017). Yet, deep tissue imaging toward applications in dermatology or gastroenterology is technically challenging due to multiple light scattering (Yu et al. 2015). There are several attempts to overcome multiple light scattering including optical coherence tomography using wavefront shaping (Jang et al. 2013; Park et al. 2018c; Yu et al. 2014; Yu et al. 2016) or accumulating single scattering light in deep tissue imaging (Kang et al. 2015).

Various research results, in which RI was utilized as imaging contrast, are highlighted in this review article. Nonetheless, we believe that there are still uncountable potential applications, which are not yet discovered. In order to fully explore the potential and capability of RI-based approaches, interdisciplinary collaborations between biologists, medical specialists, and physicists are crucial. Considering the recent rapid growth of the field and the potentials of the approach, we are optimistic that optical imaging techniques based on RI will play important roles in various topics of studies where label-free and quantitative live cell imaging is important.

## Author contributions

YKP contributed to the overall layout, figures, writing, and revision. All other authors participated in parts of the writing of this article and the survey of related literature.

